# Longitudinal samples of bacterial genomes potentially bias evolutionary analyses

**DOI:** 10.1101/103465

**Authors:** B.J. Arnold, W.P. Hanage

## Abstract

Samples of bacteria collected over a period of time are attractive for several reasons, including the ability to estimate the molecular clock rate and to detect fluctuations in allele frequencies over time. However, longitudinal datasets are occasionally used in analyses that assume samples were collected contemporaneously. Using both simulations and genomic data from *Neisseria gonorrhoeae, Streptococcus mutans, Campylobacter jejuni, and Helicobacter pylori*, we show that longitudinal samples (spanning more than a decade in real data) may suffer from considerable bias that inflates estimates of recombination and the number of rare mutations in a sample of genomic sequences. While longitudinal data are frequently accounted for using the serial coalescent, many studies use other programs or metrics, such as Tajima’s D, that are sensitive to these sampling biases and contain genomic data collected across many years. Notably, longitudinal samples from a population of constant size may exhibit evidence of exponential growth. We suggest that population genomic studies of bacteria should routinely account for temporal diversity in samples or provide evidence that longitudinal sampling bias does not affect conclusions.

## Introduction

Evolutionary analysis of bacterial genomes provides insights into the origins of diversity and is increasingly used to inform control measures for infectious pathogens. Many analyses based on simple population genetic models, such as coalescent or diffusion theory (Kingman 1982; Kimura 1964), assume individuals are sampled contemporaneously (*i.e.* from the same generation). This assumption is reasonable for data from eukaryotes, which typically have longer generation times such that samples across years may only differ by a few generations. However, bacteria have shorter generations such that a sample collected across years could violate this assumption, and the consequences for the inference of evolutionary parameters from such data have not been extensively studied. An important exception is the serial coalescent (implemented in BEAST; Drummond et al. 2002) which can account for such differences in sampling times and has been used extensively to reconstruct the history of microbes. In many cases longitudinal samples may be purposefully collected to estimate mutation rates from ‘measurably evolving populations’ (Drummond et al. 2003). Nonetheless, serial coalescent methods do not currently allow for inference of selection or complex demographic scenarios and so researchers may choose other methods with different features to analyze population genetic data, such as the popular algorithms that fit models to data using the mutation site-frequency spectrum (SFS; e.g. Excoffier et al. 2013, Gutenkunst et al. 2009). Coalescent methods may also not allow homologous recombination, which is quite common in many bacteria and may be quantified using patterns of linkage disequilibrium or phylogenetic congruence (Smith et al. 1993; Suerbaum et al. 1998; Feil et al. 2001).

We use simulations to show that longitudinal samples have an excess of rare mutations compared to contemporaneous data and can exhibit more evidence of recombination. These patterns can be seen in real genomic datasets, including previously published samples *N. gonorrhoeae, S. mutans, C. jejuni*, and *H. pylori* as examples. Our results suggest that, at least for some bacterial species, longitudinal samples spanning ~10 years have biased summary statistic values that can result in misleading demographic inference or between-species comparisons of recombination rates (e.g. Smith et al. 1993; Feil et al. 2001), especially if species differ in generation time, population size, or sample composition. Thus, researchers doing analyses sensitive to the shape of the genealogy should either account for different sampling times or provide sufficient evidence that longitudinal sampling biases do not affect conclusions.

## Results and Discussion

### Longitudinal samples have distinct genealogies

The process of coalescence describes the underlying genealogy of a sample of sequences, which dictates the patterns of genetic diversity we observe. Sequences collected from different generations (*i.e.* a longitudinal sample) cannot coalesce until their ancestral lineages are *simultaneously* present. Until this occurs (going backwards in time – the period noted *T*_*samp*_ in Figure 1), particular lineages may mutate and recombine but cannot coalesce, distorting genealogical structure. These genealogical distortions are negligible if the evolutionary time (in generations) separating longitudinal samples (*T*_*samp*_) is small in comparison to the mean time to pairwise coalescence (*T*_*coal*_) in which mutation and recombination occur in contemporaneous samples (Depaulis et al. 2009). However, any factors that decrease *T*_*coal*_, such as smaller effective population sizes from transmission bottlenecks, or increase *T*_*samp*_, such as shorter generation times (in years, the timescale on which samples are collected) may cause longitudinal samples to have distinct differences from contemporaneous ones.

**Figure 1.**
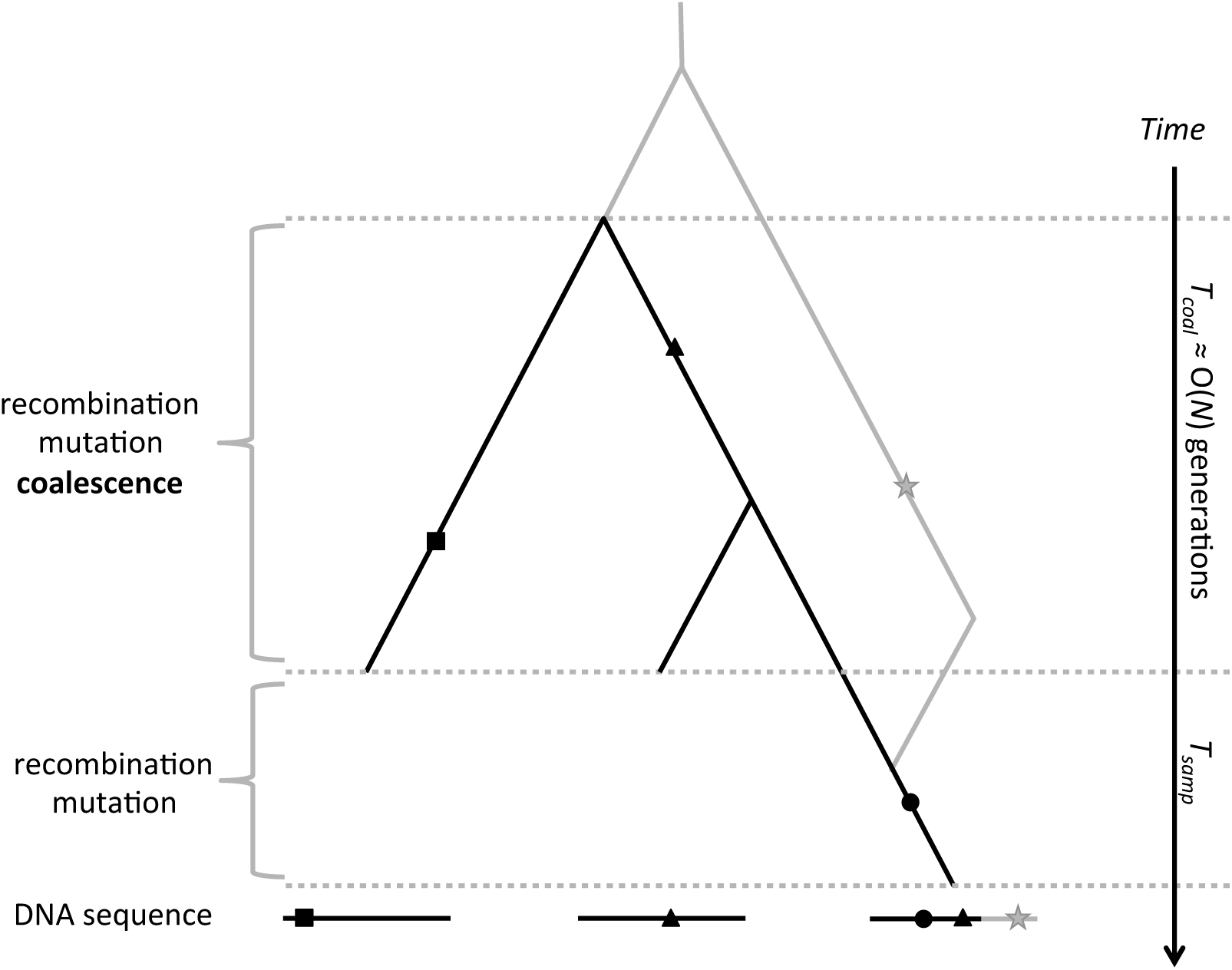
Ancestral recombination graph for a longitudinal sample. The genealogies for three DNA sequences are shown, which vary across positions due to recombination (black vs. gray lines). Shapes represent mutations. The rightmost lineage was sampled *T*_*long*_ generations later in time such that it experienced additional mutation (black circle) and recombination (bifurcation backwards in time) events. Coalescence only occurs when ancestral lineages are simultaneously present in the same population (during *T*_*coal*_) and happens over a timescale proportional to population size (*N*).

We simulated three different sampling schemes in which we longitudinally sampled sequences through time in different ways (Figure 2A). For each sampling scheme, we varied the timespan between the first and last sample, using either 0.2*N* or 0.5*N* generations, where *N* is the population size. Compared to a contemporaneous sample, longitudinal samples had SFS with an excess of rare single-nucleotide polymorphisms (SNPs; Figure 2B;Depaulis et al. 2009) and exhibited more evidence of historical recombination as measured by pairwise phylogenetic compatibility (PC) between pairs of mutations (Wilson 1965, Figure 2C). The time spanning sample collection affected observed patterns of genetic variation much more than the particular sampling schemes used here, although the asymmetric sample (scheme 3) with more lineages from certain time points was slightly less biased (Figure 2). Other unexplored sample structures, such as those with a vast majority of lineages coming from a similar generation, would likely look more like contemporaneous samples.

**Figure 2.**
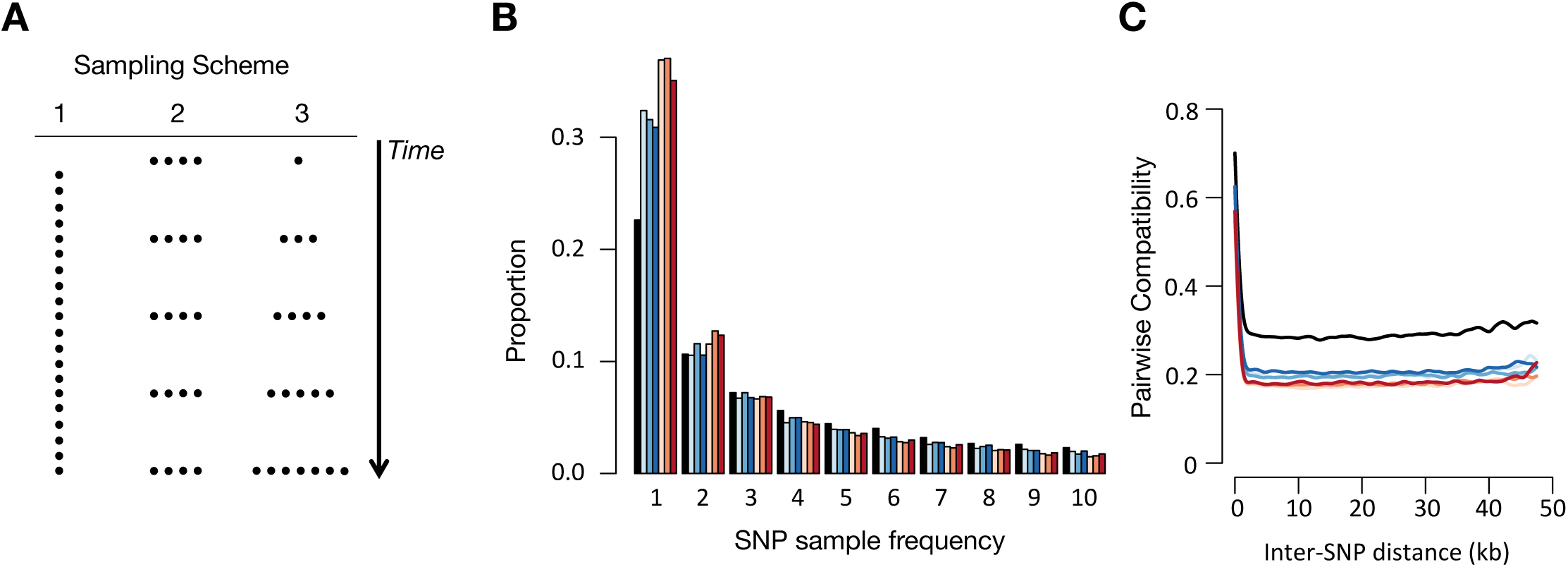
Simulated longitudinal datasets have an excess of rare variants and more evidence of recombination. **(A)** We simulated three longitudinal sampling schemes with distinct structures but with the same time spanning the first and last sample. **(B)** Longitudinal samples spanning 0.2*N* (blue) or 0.5*N* (red) generations had SFS with more rare SNPs than a contemporaneous sample (black). The particular sample structure had little effect, with light, medium, or dark blue/red corresponding to sample scheme 1, 2, or 3, respectively. **(C)** Mean pairwise compatibility between SNP pairs as a function of distance is consistently lower for longitudinal samples. Here, we simulated 50 kb fragments with a population mutation rate (2*N*μ) and recombination rate (2*Nr*) of 0.01 per site and a mean homologous recombination tract length of 1 kb. We sampled 50 individuals and calculated the mean across 20 simulations.

SFS for nonsynonymous sites under purifying selection were also more skewed by sampling bias, but overall levels of purifying selection as measured by ratios of nonsynonymous to synonymous diversity were roughly similar across all samples (Figure S1).

### Bacterial genetic samples may span relevant evolutionary timescales

To illustrate the potential for bacteria to have genealogies distorted by longitudinal sampling, we used genomic datasets from four bacterial pathogens sampled over time: *N. gonorrhoeae* (Grad et al. 2016; spanning 13 years), *S. mutans* (Cornejo et al. 2013; spanning 27 years), *C. jejuni* (Sheppard et al. 2013; Yahara et al. 2014; spanning 11 years), and *H. pylori* (Blanchard et al. 2013; spanning 11 years). We selected these datasets because they not only contained longitudinal samples spanning more than a decade from a restricted geographic location but also had enough samples from a given year for meaningful comparison (i.e. greater than or equal to 10). Population structure may also frequently create biases in evolutionary inference, but this has been studied elsewhere (e.g. Lapierre et al. 2016), so we attempted to minimize the effects of structure by only looking at samples within a city (San Diego, California, USA for *N. gonorrhoeae* or Cleveland, Ohio, USA for *H. pylori*), country (United Kingdom for *S. mutans*), or sequence type (ST45 for *C. jejuni*).

For three species examined, a sufficient number of generations separate longitudinal samples (*T*_*long*_) with respect to population size (*T*_*coal*_) to distort genealogies and create measurable differences between summary statistics: serial samples exhibit more evidence of historical recombination (although only slightly for *C. jejuni*) and have skewed SFS with an excess of rare mutations (Figure 3B). SFS for larger samples, which are unavoidably longitudinal, are either similar or slightly more skewed than subsamples (Table S1). These summary statistic biases could thus affect within-species estimation of recombination rates (e.g. Takuno et al. 2012) or between-species comparisons (e.g. Smith et al. 1993; Feil et al. 2001) if sampling dates are not accounted for, especially if datasets differ in sampling timespans or species have different generation times or effective population sizes (e.g. Suerbaum et al. 1998 which compares bacteria with a eukaryote). These biases could also create problems for any application that relies on the shape of the SFS, such as the calculation of Tajima’s D (as in Touchon et al. 2014) or programs that use the SFS to fit demographic models to data, such as δaδi, *prfreq,* or *fastsimcoal2* (used in Cornejo et al. 2013; Pepperell et al. 2013; Montano et al. 2015, respectively). However, these biases could be favorable for other applications such as genome-wide association studies since longitudinal samples would exhibit less linkage than contemporaneous ones.

**Figure 3.**
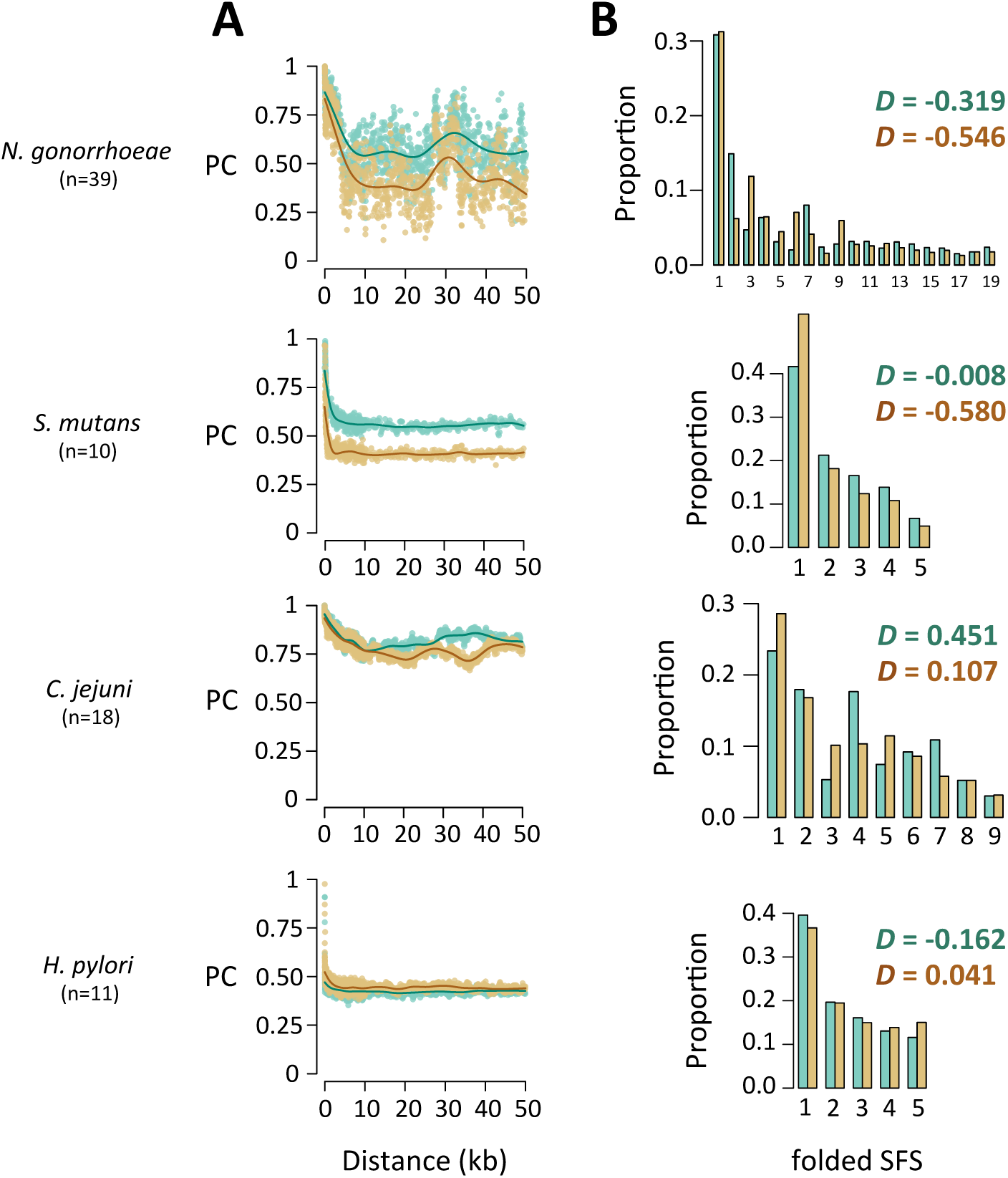
Longitudinal samples in real bacterial datasets span relevant timescales. Pairwise compatibility (PC) as a function of inter-SNP distance **(A)** and the SFS along with Tajima’s D **(B)** are shown for contemporaneous (green) and longitudinal (brown) samples of the four species analyzed in this study. Sample sizes used in analyses are shown below the species name.

Summaries for longitudinal and contemporaneous *H. pylori* samples appear quite similar (Figure 3). Potential explanations include larger population sizes, longer generation times, or other factors such as population structure.

To illustrate, we used *fastsimcoal2* (Excoffier et al. 2013) to fit two demographic models, either constant population size or exponential growth, to all samples. In agreement with the sample SFS in Figure 3B, model fits to longitudinal samples either had grossly inflated signals of population growth for *N. gonorrhoeae* or, for *S. mutans*, provided evidence for growth when contemporaneous samples were better explained by a model of constant size (Table 1, Figure S2). The 8-fold population growth estimated from the longitudinal *S. mutans* sample is similar to that reported to the 5-fold growth in Cornejo et al. 2013. Thus, while these results are not definitive, they still provide evidence that longitudinal sampling bias may have contributed to some, if not all, of the signal of growth in *S. mutans*. Masking rare variants (e.g. singletons and doubletons) does not ameliorate these biases, according to growth model fits to simulated datasets with larger sample sizes (n=50, Table S2).

**Table 1.** Maximum likelihood estimates of exponential growth model parameters

| Species | Contemporaneous sample |  | Longitudinal sample |  |
| --- | --- | --- | --- | --- |
| | Fold growth<br>($N_C/N_A$) | Time of growth<br>($N_A$ generations) | Fold growth<br>($N_C/N_A$) | Time of growth<br>($N_A$ generations) |
| *N. gonorrhoeae* | 4.32 | 0.08 | 34.25 | 0.02 |
| *S. mutans* | NA* | NA* | 8.12 | 0.24 |

Notes: The exponential growth model involves an ancestral population of size $N_A$ that starts growing at some point in time to contemporary size $N_C$ . Models were fit to the SFS of fourfold-degenerate sites.
\*A model of constant population size fit the data better than a model of exponential growth.

## Conclusions

Samples collected over time are common in the growing literature on the population genomics of bacteria. This reflects analyses of samples already collected and in freezers, but also a deliberate strategy to examine the way populations change over time. We have found that longitudinal sample schemes can produce erroneous signals of population growth and exaggerated rates of recombination if sample dates are ignored. This happens for intuitive reasons illustrated in Figure 1; the longer sampling period provides more generations for mutation or recombination to occur, skewing the SFS and the total amount of observed recombination. This can generate wholly artificial signals of population growth.

While our results urge caution in interpreting evolutionary analyses when collection dates are not accounted for, this problem is not expected to affect species with small *T*_*long*_ (longer generations or near-contemporaneous samples) and/or large *T*_*coal*_ (large population sizes), such as non-pathogenic bacteria that have less population structure and do not experience frequent population bottlenecks from limited transmission. However, bacterial genomic samples frequently span more than a decade and may have significant biases. We thus suggest that sampling dates and proof that analyses do not suffer from longitudinal sampling bias should be routinely provided in evolutionary genetic studies of bacteria.

## Methods

### Bacterial Genomic Data

*N. gonorrhoeae* data were kindly provided by Yonatan Grad (Grad et al. 2016), *C. jejuni* data were provided by Samuel Sheppard (Sheppard et al. 2013, Yahara et al. 2014), and we downloaded *S. mutans* data used in Cornejo et al. 2013 and *H. pylori* data reported in Blanchard et al. 2013 from NCBI. We analyzed *de novo* assemblies with PROKKA (Seemann 2014), using an amino acid file from a reference genome (FA1090 for *N. gonorrhoeae*, UA159 for *S. mutans*, CjeNCTC11168 for *C. jejuni*, and HPY26695 for *H. pylori*). We then identified core and accessory genes with ROARY (Page et al. 2015) and used only core genes that were also present in the reference genome for analysis of PC and the SFS. We inferred position information between polymorphic sites using the relative positions of genes in the reference genome, not from a reference-based DNA alignment. We calculated PC (Wilson 1965) and Tajima’s D (Tajima 1989) form core gene alignments using custom Perl scripts. All contemporaneous and serial samples used in this study may be found in Table S3.

### Simulations of longitudinal datasets

We designed a forward-in-time simulator of a Wright-Fisher population in C++. We used this program to simulate a population of *N*=1000 haploid genomes for a burn-in time of 10*N* generations until the population reached mutation-drift balance. After this time, we either took a contemporaneous or longitudinal sample of n=50 genomes. Longitudinal samples spanned either 200 (0.2*N*) or 500 (0.5*N*) generations between the first and last sample, and we used three different sampling schemes (Figure 2A). For scheme one, we sampled 1 genome every 4 or 10 generations until n=50, such that samples were collected across ~0.2*N* or ~0.5*N* generations, respectively. Likewise, for scheme two, we sampled 10 genomes every 50 or 125 generations until n=50. Lastly, for scheme three we took five samples of size 18, 14, 10, 6, and 2 that were separated by 50 or 125 generations. We note that while real bacterial populations sizes are likely much larger than the size simulated here, our results scale to populations of arbitrary size as long as time is measured in *N* generations. For the results in Figure 2, we simulated 50 kb fragments and fewer repetitions (20), but for the analyses of purifying selection in Figure S1, we simulated 10 kb fragments and more repetitions (1000).

### Demographic model fitting

We fit both a model of constant population size and exponential growth to the SFS of fourfold degenerate sites using *fastsimcoal2* (Excoffier et al. 2013). For each sample SFS, we ran *fastsimcoal2* 50 times, which uses an expectation-maximization algorithm to search parameter space. We chose the run that produced the highest likelihood for model selection, and we used Akaike information criterion (AIC) to evaluate which model had the higher probability of being correct given the candidate set of models (Figure S2). To explore the effect of masking rare variants for parameter estimation, we fit exponential growth models to simulated datasets using either the full SFS (default) or a minimum mutation count of three (by including the “-C 3” flag in the *fastsimcoal2* command; Table S2).

## Acknowledgements

We would like to thank Marc Lipsitch, Hsiao-Han Chang, and Omar Cornejo for useful discussion. We would also like to thank Michael Stanhope for providing the sampling dates of the *S. mutans* isolates.

---

**Figure S1.**
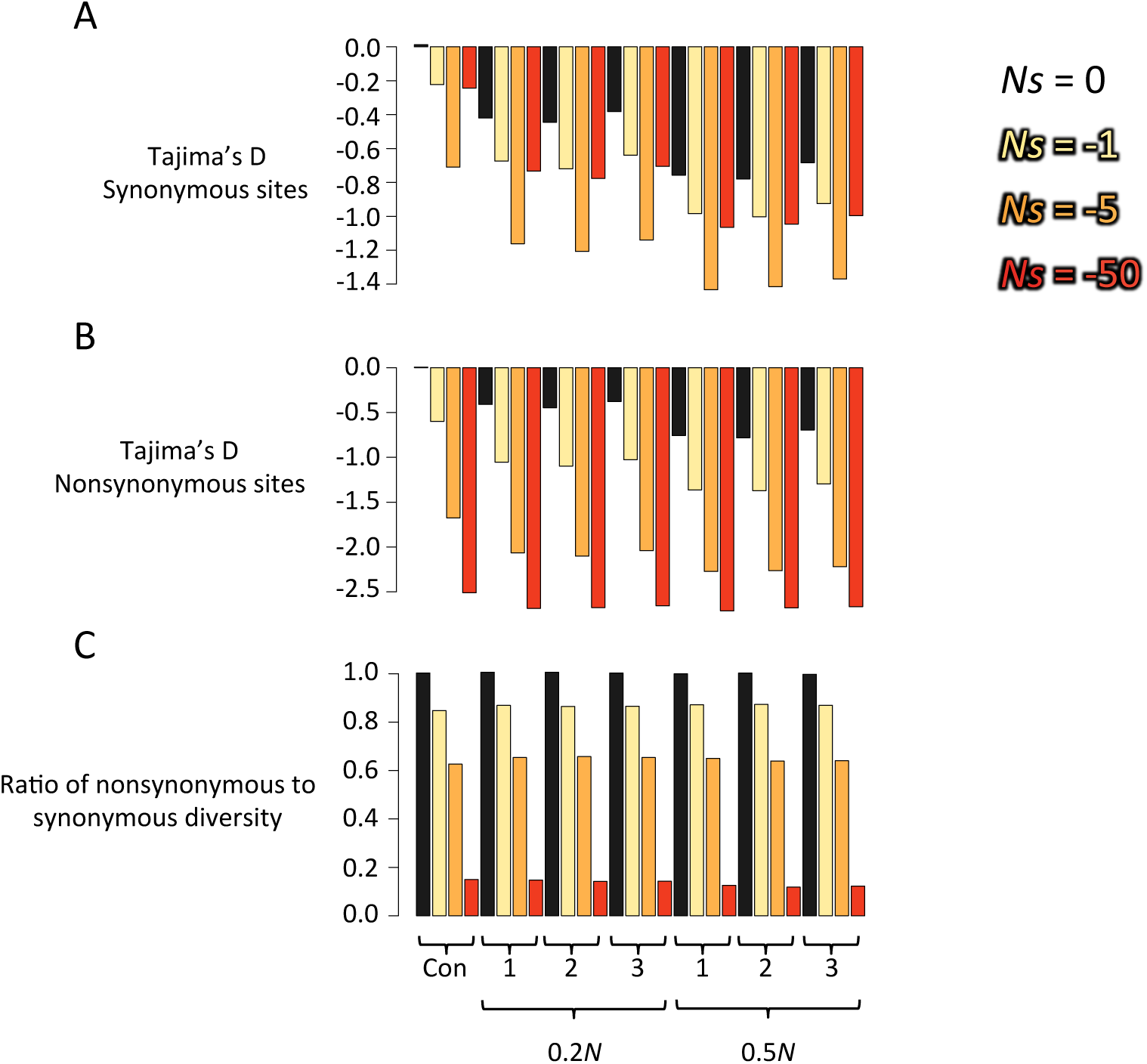
Analysis of synonymous and nonsynonyous variation for longitudinal samples. Shown are summary statistics from simulation studies of a contemporaneous sample (Con) and various longitudinal samples spanning either 0.2*N* or 0.5*N* generations and having sample structures according to those shown in Figure 2A (1, 2, or 3). We simulated samples collected from a neutral population (*Ns*=0) or with varying degrees of purifying selection (*Ns*=-1, −5, or −50). We calculated Tajima’s D at both synonymous (**A**) and nonsynonymous (**B**) sites and the ratio of nonsynonymous to synonymous diversity (**C**, calculated as Watterson’s estimator). Tajima’s D becomes more negative at synonymous and nonsynonymous sites with longitudinal sampling bias and from purifying selection on nonsynonymous mutations with linkage to synonymous mutations. Levels of purifying selection, as measured by ratios of nonsynonymous to synonymous diversity (**C**), do not significantly differ between contemporaneous and longitudinal samples. Here, we simulated 10 kb fragments with a population mutation rate (2*N*μ) and recombination rate (2*Nr*) of 0.01 per site and a mean homologous recombination tract length of 1 kb. We sampled 50 individuals and calculated the mean across 1000 simulations.

**Figure S2.**
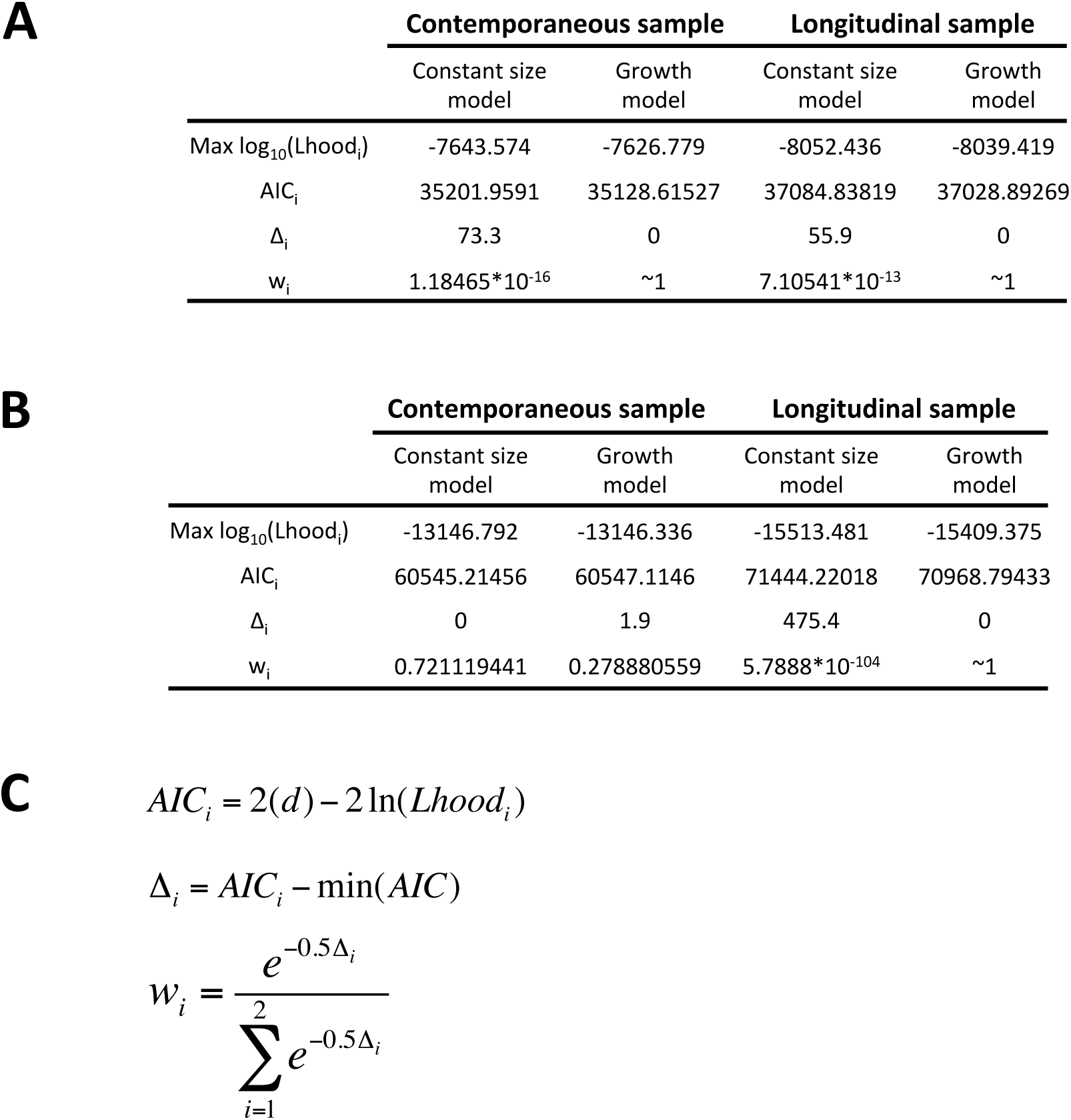
Model selection using *fastsimcoal2* and AIC. AIC analyses are shown for *N. gonorrhoeae* **(A)** and *S. mutans* **(B)** samples. For each analysis, the maximum likelihood of 50 *fastsimcoal2* runs is shown (Max log_10_(Lhood)), along with the components necessary for the calculating the Akaike weight (*w*_*i*_) for each model **(C)**, which may be interpreted as the probability that model *i* is the correct one. Here, *d* represents the number of model parameters used, which was either 1 for the constant size model or 3 for the growth model, since both an ancestral and contemporary population need to be specified, along with the time at which population growth starts.

**Table S1.** Tajima’s D for contemporaneous and longitudinal samples

| Species | Sample ID | Tajima's D* | Sample Size | Sample Description |
| --- | --- | --- | --- | --- |
| ***N. gonorrhoeae*** |  |  |  |  |
|  | sample 1 | -0.319 | 39 | Contemporaneous sample from same year (2009) and city (San Diego, California) |
|  | sample 2 | -0.546 | 39 | Serial sample from San Diego of same size as contemporaneous sample |
|  | sample 3 | -0.734 | 152 | All available samples from San Diego |
| ***S. mutans*** |  |  |  |  |
|  | sample 1 | -0.008 | 10 | Contemporaneous sample from same year (2006) and country (UK) |
|  | sample 2 | -0.579 | 10 | Serial sample from UK of same size as contemporaneous sample |
|  | sample 3 | -0.616 | 21 | All available samples from UK |
| ***C. jejuni*** |  |  |  |  |
|  | sample 1 | 0.451 | 18 | Contemporaneous sample from same year (2008) and sequence type (ST45) |
|  | sample 2 | 0.107 | 18 | Serial sample of ST45 of same size contemporaneous sample |
|  | sample 3 | -0.383 | 52 | All available samples from ST45 |
| ***H. pylori*** |  |  |  |  |
|  | sample 1 | -0.162 | 11 | Contemporaneous sample from same year (1986) and city (Cleveland, Ohio) |
|  | sample 2 | 0.041 | 11 | Serial sample from Cleveland of same size as contemporaneous sample |
|  | sample 3 | -0.315 | 38 | All available samples with dates from Cleveland |

\*Calculated from the SFS of fourfold degenerate sites genome-wide

**Table S2.** Parameter estimates for growth model are similar when rare variants are masked

| Sampling scheme | Sample spanning 0.2N generations |  |  |  | Sample spanning 0.5N generations |  |  |  |
| --- | --- | --- | --- | --- | --- | --- | --- | --- |
|  | Full SFS |  | Minimum mutation count of 3 |  | Full SFS |  | Minimum mutation count of 3 |  |
|  | Fold growth | Time of growth | Fold growth | Time of growth | Fold growth | Time of growth | Fold growth | Time of growth |
| 1 | 3.79 | 0.11 | 3.66 | 0.11 | 6.28 | 0.13 | 6.46 | 0.13 |
| 2 | 3.22 | 0.12 | 3.27 | 0.11 | 5.23 | 0.19 | 5.28 | 0.19 |
| 3 | 2.87 | 0.11 | 2.91 | 0.11 | 4.65 | 0.15 | 4.77 | 0.14 |

Notes: The exponential growth model involves an ancestral population of size $N_A$ that starts growing at some point in time to contemporary size $N_C$ . Fold growth was measured as $N_C/N_A$ , and time of growth is measured in $N_A$ generations. Growth models were fit as described in Methods.

**Table S3.** Collection dates of strains used in contemporaneous and longitudinal samples.

| Contemporaneous sample |  | Longitudinal sample |  |
| --- | --- | --- | --- |
| Strain name | Collection Date | Strain name | Collection date |
| 8289_2#74 | 2009 | 17150_8#82 | 2000 |
| 8727_5#56 | 2009 | 17176_1#85 | 2001 |
| 8289_2#68 | 2009 | 17225_3#34 | 2002 |
| 8289_2#20 | 2009 | 17150_8#17 | 2002 |
| 8727_5#58 | 2009 | 17150_8#21 | 2002 |
| 8289_2#72 | 2009 | 17150_8#19 | 2002 |
| 8727_5#68 | 2009 | 17150_8#32 | 2003 |
| 8727_5#48 | 2009 | 17150_8#58 | 2004 |
| 8727_5#74 | 2009 | 17225_2#15 | 2004 |
| 9716_3#90 | 2009 | 17225_2#14 | 2004 |
| 8727_8#24 | 2009 | 17150_8#61 | 2004 |
| 8289_2#76 | 2009 | 15335_4#28 | 2005 |
| 8727_8#66 | 2009 | 15335_4#29 | 2005 |
| 15335_5#85 | 2009 | 15335_4#30 | 2005 |
| 8727_8#20 | 2009 | 15335_5#24 | 2005 |
| 8289_2#28 | 2009 | 15335_5#66 | 2007 |
| 8727_5#70 | 2009 | 15335_4#69 | 2007 |
| 8727_8#18 | 2009 | 15335_5#67 | 2007 |
| 8727_8#64 | 2009 | 15335_6#1 | 2007 |
| 8727_5#66 | 2009 | 8289_2#74 | 2009 |
| 8289_2#30 | 2009 | 8727_5#56 | 2009 |
| 8727_8#54 | 2009 | 8289_2#68 | 2009 |
| 8289_2#78 | 2009 | 15335_5#85 | 2009 |
| 8727_5#64 | 2009 | 8727_5#54 | 2010 |
| 8727_8#34 | 2009 | 8727_5#43 | 2010 |
| 8727_8#38 | 2009 | 8727_5#45 | 2010 |
| 8727_5#62 | 2009 | 8727_8#96 | 2010 |
| 8727_5#52 | 2009 | 15335_6#16 | 2011 |
| 8289_2#70 | 2009 | 15335_3#74 | 2011 |
| 8289_2#32 | 2009 | 15335_6#67 | 2011 |
| 8727_8#36 | 2009 | 15335_4#93 | 2011 |
| 8289_2#22 | 2009 | 15335_2#60 | 2012 |
| 8289_2#26 | 2009 | 16043_2#23 | 2012 |
| 8289_2#67 | 2009 | 15335_2#2 | 2012 |
| 8289_2#69 | 2009 | 15335_2#64 | 2012 |
| 8289_2#71 | 2009 | 15335_2#75 | 2013 |
| 8289_2#73 | 2009 | 15335_2#68 | 2013 |
| 8289_2#75 | 2009 | 15335_7#67 | 2013 |
| 8289_2#77 | 2009 | 15335_7#1 | 2013 |

***S. mutans***
| Contemporaneous sample |  | Longitudinal sample |  |
| --- | --- | --- | --- |
| Strain name | Collection Date | Strain name | Collection date |
| A19 | 2006 | N3209 | 1979 |
| A9 | 2006 | N29 | 1991 |
| G123 | 2006 | NFSM1 | 1994 |
| M21 | 2006 | N34 | 1997 |
| M2A | 2006 | N66 | 1997 |
| ST1 | 2006 | NLML1 | 2001 |
| ST6 | 2006 | NLML4 | 2001 |
| T4 | 2006 | NLML8 | 2002 |
| U138 | 2006 | A19 | 2006 |
| W6 | 2006 | A9 | 2006 |

***C. jejuni***
| Contemporaneous sample |  | Longitudinal sample |  |
| --- | --- | --- | --- |
| Strain name | Collection Date | Strain name | Collection date |
| 1702 | 2008 | 112 | 2004 |
| 1722 | 2008 | 114 | 2004 |
| 1723 | 2008 | 1775 | 2005 |
| 1727 | 2008 | 1776 | 2005 |
| 1733 | 2008 | 39 | 2005 |
| 1737 | 2008 | 1771 | 2006 |
| 1738 | 2008 | 45 | 2006 |
| 1740 | 2008 | 1800 | 2007 |
| 288 | 2008 | 1817 | 2007 |
| 309 | 2008 | 405 | 2007 |
| 312 | 2008 | 1702 | 2008 |
| 321 | 2008 | 1722 | 2008 |
| 324 | 2008 | 1723 | 2008 |
| 378 | 2008 | 342 | 2009 |
| 406 | 2008 | 371 | 2009 |
| 416 | 2008 | 382 | 2009 |
| 429 | 2008 | 434 | 2011 |
| 472 | 2008 | 5006 | 2015 |

***H. pylori***
| Contemporaneous sample |  | Longitudinal sample |  |
| --- | --- | --- | --- |
| Strain name | Collection Date | Strain name | Collection date |
| HpA_14 | 6/6/1986 | HpA_8 | 4/28/1986 |
| HpA_16 | 6/18/1986 | HpA_27 | 12/1/1986 |
| HpA_17 | 6/18/1986 | HpP_11 | 6/9/1987 |
| HpA_20 | 10/10/1986 | HpP_16 | 12/17/1987 |
| HpA_26 | 12/1/1986 | HpH_4 | 9/20/88 |
| HpA_27 | 12/1/1986 | HpH_27 | 3/23/1989 |
| HpA_4 | 4/18/1986 | HpP_26 | 11/9/1989 |
| HpA_5 | 4/23/1986 | HpP_30 | 6/28/1990 |
| HpA_6 | 5/2/1986 | HpP_41 | 4/12/1991 |
| HpA_8 | 4/28/1986 | HpP_62 | 3/13/1996 |
| HpP_8 | 4/28/1986 | HpP_74 | 3/6/1997 |

## Citations

Blanchard, T.G. et al., 2013. Genome sequences of 65 helicobacter pylori strains isolated from asymptomatic individuals and patients with gastric cancer, peptic ulcer disease, or gastritis. Pathogens and Disease, 68(2), pp.39–43.

Cornejo, O.E. et al., 2013. Evolutionary and population genomics of the cavity causing bacteria streptococcus mutans. Molecular Biology and Evolution, 30(4), pp.881–893.

Depaulis, F., Orlando, L. & Hänni, C., 2009. Using classical population genetics tools with heterochroneous data: Time matters! PLoS ONE, 4(5).

Drummond, A.J. et al., 2002. Estimating mutation parameters, population history and genealogy simultaneously from temporally spaced sequence data. Genetics, 161(3), pp.1307–1320.

Drummond, A.J. et al., 2003. Measurably evolving populations. Trends in Ecology and Evolution, 18(9), pp.481–488.

Excoffier, L. et al., 2013. Robust Demographic Inference from Genomic and SNP Data. PLoS Genetics, 9.

Feil, E.J. et al., 2001. Recombination within natural populations of pathogenic bacteria: short-term empirical estimates and long-term phylogenetic consequences. Proceedings of the National Academy of Sciences of the United States of America, 98(1), pp.182–187.

Grad, Y.H. et al., 2016. Genomic epidemiology of gonococcal resistance to extended-spectrum cephalosporins, macrolides, and fluoroquinolones in the United States, 2000-2013. J Infect Dis, 214(10), pp.1579–87.

Gutenkunst, R.N. et al., 2009. Inferring the joint demographic history of multiple populations from multidimensional SNP frequency data. PLoS Genetics, 5(10).

Kimura, M., 1964. Diffusion Models in Population Genetics. Journal of Applied Probability, 1(2), pp.177–232.

Kingman, J.F.C., 1982. On the Genealogy of Large Populations. Journal of Applied Probability, 19, p.27.

Lapierre, M. et al., 2016. The impact of selection, gene conversion, and biased sampling on the assessment of microbial demography., pp.1–15.

Montano, V. et al., 2015. Worldwide Population Structure, Long Term Demography, and Local Adaptation of Helicobacter pylori. Genetics, 200(July), pp.947–963.

Page, A.J. et al., 2015. Roary: Rapid large-scale prokaryote pan genome analysis. Bioinformatics (Oxford, England), pp.1–3.

Pepperell, C.S. et al., 2013. The Role of Selection in Shaping Diversity of Natural M. tuberculosis Populations. PLoS Pathogens, 9(8).

Seemann, T., 2014. Prokka: rapid prokaryotic genome annotation. Bioinformatics, 30(14), pp.2068–2069.

Sheppard, S.K. et al., 2013. Genome-wide association study identifies vitamin B 5 biosynthesis as a host specificity factor in Campylobacter. Proceedings of the National Academy of Sciences of the United States of America, 110(May), pp.1–5.

Smith, J.M. et al., 1993. How clonal are bacteria? Proceedings of the National Academy of Sciences of the United States of America, 90(10), pp.4384–4388.

Suerbaum, S. et al., 1998. Free recombination within Helicobacter pylori. PNAS, 95(October), pp.12619–12624.

Tajima, F., 1989. Statistical method for testing the neutral mutation hypothesis by DNA polymorphism. Genetics, 123, pp.585–595.

Takuno, S. et al., 2012. Population genomics in bacteria: A case study of Staphylococcus aureus. Molecular Biology and Evolution, 29(2), pp.797–809.

Touchon, M. et al., 2014. The genomic diversification of the whole Acinetobacter genus: origins, mechanisms, and consequences. Genome biology and evolution, 6(10), pp.2866–82.

Wilson, E.O., 1965. A Consistency Test for Phylogenies Based on Contemporaneous Species. Systematic Zoology, 14(3), pp.214–220.

Yahara, K. et al., 2014. Efficient inference of recombination hot regions in bacterial genomes. Molecular Biology and Evolution, 31(6), pp.1593–1605.

